# The reconstitution of body mass index in HIV positive subjects under antiretroviral treatment in Kinshasa

**DOI:** 10.1101/524462

**Authors:** Guyguy Kabundi Tshima, Paul Madishala Mulumba

## Abstract

**Objective:** We aimed to evaluate BMI changes in HIV adults’ subjects in the first year of ART in malaria endemic areas.

**Methods:** We used linear regression analysis showing that the change in weight at 12 months (y) in a malaria-endemic area is related to malaria infection at admission and its different episodes as illustrated by equation: y = a + bxi + ε, where x is malaria on admission, i refers to episodes of clinical malaria infection during the year, b is the slope, a is a constant and ε are confounding factors such as tuberculosis or poor eating habits.

**Results:** We found a positive value for b (b = 0.697), and this shows that weight loss at 12 months is correlated with the diagnosis of severe malaria at admission. In other words, severe malaria eliminates the weight gained under ART.

**Conclusions:**

1. Malaria is the leading cause of weight loss under ART.
2. Important recommendation for future:

This study suggests nutritional education based on local foods containing antioxidants to fight the oxidative stress generated by HIV and stimulated by *Plasmodium falciparum* during febrile episodes. Oxidative stress is blocked by NADPHase which is a metalloenzyme based on selenium.

Thus, to prevent a weight loss or the occurrence of the protein-energy malnutrition among people living with HIV, it is necessary to use the nutritional education.

**Résumé:** *Objectif:* Nous voulions évaluer les modifications de l’IMC chez les patients VIH adultes au cours de la première année du traitement antirétroviral dans une zone d’endémie palustre

*Matériel et Méthodes:* Nous avons utilisé une analyse de régression linéaire montrant que la variation de poids à 12 mois (y) dans une zone d’endémie palustre est liée à l’infection palustre à l’admission et à ses différents épisodes, comme l’illustre l’équation suivante: y = a + bxi + ε, où x est le paludisme à l’admission, i les épisodes de paludisme clinique survenus au cours de l’année, b est la pente, a est une constante et ε sont des facteurs de confusion tels que la tuberculose ou de mauvaises habitudes alimentaires..

*Résultats:* Nous avons trouvé une valeur positive pour b (b = 0,697), ce qui montre que la perte de poids à 12 mois est en corrélation avec le diagnostic de paludisme grave à l’admission. En d’autres termes, le paludisme grave élimine le poids gagné sous traitement antirétroviral.

*Conclusions:*

1. Le paludisme est la principale cause de perte de poids sous ARV.
2. Recommandation importante pour l’avenir : Cette étude suggère une éducation nutritionnelle basée sur des aliments locaux contenant des anti-oxydants pour lutter contre le stress oxydatif généré par le VIH et stimulé par le *Plasmodium falciparum* lors des poussées fébriles. Le stress oxydatif est bloqué par la NADPHase qui est une métalloenzyme à base de sélénium. Ainsi, il est nécessaire d’utiliser l’éducation nutritionnelle pour prévenir la perte du poids sous ARV.

## Introduction

For the investigation on weight evolution under ART in case of malaria being reported, worldwide in 2016, there were 36.7 million people living with HIV [1]. And in 2015, there were an estimated 214 million cases of malaria worldwide, and an estimated 438 000 deaths [1]. Approximately 90% of all malaria deaths occur in Africa [2]. Since 2011 there was no WHO recommendation regarding any specific antimalarial treatment (AMT) for patients living with HIV in malaria areas [3], there is a need to explore the possibility of establishing specific guidelines for this category of patients co-infected with *Plasmodium falciparum*. The present work deals with the pending situation of co-infection HIV-malaria which remains a major public health problem in several countries worldwide [3].

A first step towards the physiopathology of the weight loss during the co-infection severe malaria-HIV because due to co-infection, the metabolic demands of antioxidant products such as selenium, vitamin C and vitamin E are increased. As a result, micronutrient deficiencies increase due to malaria and HIV [4].

The mechanisms of HIV oxidative stress and malaria progression can be explained in two ways: first, malaria stimulates NADPHase blocked by antiretroviral therapy, exposing the patient to weight loss, and elsewhere it has been shown that HIV-positive people and those infected with malaria suffer from an oxidative / antioxidant imbalance [5]. Therefore, establishing the physiological balance between oxidants and antioxidant factors is of great therapeutic interest. In this context, the implication of the present study is to evaluate the impact of severe malaria on weight change and the perspective is to design a nutrient (Selenium) that can help strengthen the immune state of HIV+ subjects to fight HIV-induced oxidative stress and *Plasmodium falciparum* during severe AIDS-malaria co-infection [6].

We expected dietary intake using local foods rich in antioxidants. A first step towards our research began with foods inquiry when in 2007, according to an American work, the daily intake of Selenium in the form of food supplements could even reduce the viral load in patients with HIV: a study of 262 patients, the antioxidant properties of Selenium would be responsible for this decrease. “An explanation that required confirmation,” said the author [7]. Anyway, it was an interesting thought trail. “Selenium supplementation is a simple, safe and inexpensive approach” [7]. Not to mention that to refill Selenium, there are other solutions than the use of food supplements e.g. Seafood, Mushrooms. It would thus protect against cardiovascular diseases but, also, against certain digestive cancers [7].

Several international works lend it interesting properties:

- Several investigators have found that HIV-infected patients have a compromised antioxidant defense system. Blood antioxidants are decreased, and the products of lipid and protein oxidation are increased in these patients. This may have physio pathological implications [8];
- Selenium supplementation affects specific populations of T lymphocytes and decreases the markers of lipoperoxides [9];
- Selenium plays an important role in the maintenance of immune function and neutralizes the superoxide ions produced by activated macrophages and neutrophils in response to the aggression of the body by microorganisms [10].

So, the progression of HIV infection to the AIDS stage is due to the production of free radicals. It would be interesting to identify possible mechanisms and clinical trials to evaluate the effect of Selenium supplementation in the progression of HIV infection. The oxidative stress causes the production of cytokines that lead to cachexia [11].

This study focuses on the reconstitution of weight in HIV positive patients under antiretroviral treatment (ART) in the environment of Kinshasa where patients living with HIV are combining the antiretroviral treatment (ART) and the antimalarial treatment (AMT) when they are diagnosed malaria positive with microscopy. The rationale for this study is to describe why they lose weight with a good observance on ART. What is the exposure factor?

## Methods and findings

### Data collection

We obtained weight, the individual CD_4_ count and the diagnosis of severe malaria among adults in the Democratic Republic of the Congo using the 2007 AMOCONGO Kinshasa-Kasavubu Register HIV database. The AMOCONGO Kinshasa-Kasavubu 2007 database is a routinely collected health data which a product of the daily operations of the healthcare centre are collected independently of specific a priori research questions. The use of health data routinely collected in a prospective view is explained in those studies following the RECORD-PE [12–30]

#### Ethical issues

The University of Kinshasa ethic committee and the National Programme of AIDS estimated ethic to use the health data sampling in AMOCONGO Kinshasa-Kasavubu Register.

### BMI

BMI is calculated using weight (kg) divided by squared height (m^2^). Weight and height were directly measured by AMOCONGO medical staff. Since our targeted samples are adults, we do not expect any significant variation in the height for twelve months that could affect the BMI. Therefore, only the change in weight is considered as a parameter that can be evaluated in the evolution of BMI. So, we replaced BMI by weight. Weight was reported as usual at the admission, at 3 months, at 6 months and at 12 months.

### Adequate logistic regression model

Using Minitab software, we compute the binary logistic regression after the regression, confusion, and interaction assumptions. The probabilities greater than 5% means that the model is adequate.

### Limitations

The following were limitations:

- Malaria access: only 1 or repeated access (HIV infection potentiates the frequency of access), it was necessary to distinguish between severe malaria and simple malaria
- The different ART regimes: Triomune-30, Triomune-40, Kaletra, … It is known that certain molecules are leading to more resistance than others.
- The different antimalarial drugs used and antimalarial combinations: Quinine, Sulfadoxine-Pyrimethamine, Artesunate-Amodiaquine, Arthemether-Lumefantrine, …
- Nutritional education: distinguish between patients who have attended a nutrition education course and others who have not.
- Antibiotherapy: duration and frequency of treatment
- The date of last taking deworming medication: undernutrition can be controlled by intestinal worms (Ascaris, Anguillules, Trichocephales, …). It would be interesting to note whether a stool examination was done or not
- Alcohol: plays an important role in the accumulation of fats
- Tobacco: makes you lose weight; distinguish between smokers and non-smokers
- HIV serologic status: consider the control group (HIV-)
- Marital status: married couples can have a regular diet compared to singles)
- ART duration: 3 months were sufficient to evaluate the recovery of the body mass index?
- CD4 lymphocyte count: broadly divided into two groups

* Normal CD_4_ level ≥ 200

* Low CD_4_ rate ≤ 200: those are put on ART

- Associated opportunistic pathologies

- Social standing
- Age: Adults, the ideal would be to resort to children in vaccination period (0-5 years) to limit the confounding factors
- Number of CD_4_ count for HIV-negative adults: we did not have data on the enumeration of CD_4_ lymphocytes for HIV-negative adults.

## Results

### Descriptive data

#### Demographic characteristics of study participants

Table 1 shows baseline characteristics of the Congolese adults.

**Table 1.** Sample characteristics, age distribution.

| Age in years (n = 72) | Percentage | Mean (SD) |
| --- | --- | --- |
| 21 < 29 years (n=22) | 30.7% | 32.6 (11.6) |
| 30-49 years (n =23) | 32.1% |  |
| 50-59 years (n=27) | 37.2% |  |
| ≥60 years (n = 0) | 0% |  |

The final sample included 72 individuals, which expanded to 72 adults with 22 young adults 21 to 29 years old, in comparison to adults of 30 to 49 years old, of 50 to 59 years old. The average age was 32.6 years (11.6). 37.2% (50-59 years) was the highest percentage; this means that experienced people are the most infected with HIV-AIDS. This is really a problem for a developing country like the DRC that needs experienced adult workers for its development.

Table 2 shows that the sex ratio was 4 women to 1 man. Apart from the biological reasons that would explain that the female genital area is larger than the male one and explain the vulnerability of the latter in unprotected sex. It is meanly meaning that Congolese women seek more help from health facilities than men for cultural reasons. Men want to see themselves strong and therefore not vulnerable to disease. Sick Guards are women most of the time, and for prenatal or pre-school clinics, women are more likely to attend the hospital than men. Congolese men refuse to ask for help in the first symptoms of the disease, they first want to fight alone so, logically with a weakened body the expected mortality rate should be higher in men than in women because they will be treated later, with the higher risk of death following the natural course of the disease.

**Table 2.** Repartition of sex.

| Sex (n =72) | Percentage (n =72) |
| --- | --- |
| Men (n =15) | 21% (n/n ) |
| Women (n=57) | 79% (n/n ) |

### Significance of logistic regression model after analysis of health data

Table 3 showed a p-value> 0.05, at this level we cannot draw a conclusion because it has no scientific value, so we did linear regression to try to show a correlation between the effect of severe malaria on the number of CD_4_ and the decrease in weight. Due to the binary logistic regression a positive correlation was found between the number of CD_4_ <50 cells / μl and severe malaria **on** admission, but not significantly (p-value> 0.05).

**Table 3.** Multivariate logistic regression analysis of clinical malaria infection among 72 HIV patients living in malaria endemic area and their precision (eg, 95% confidence interval).

| Severe malaria | OR (95%CI) | *P*- value |
| --- | --- | --- |
| Number CD<sub>4</sub> < 50 cells / µl) | 1.19 (0.19-7.56) | 0.854 |
| Initial Weight | 0.86 (0.71-1.04) | 0.113 |
| Weight at 12 months | 1.10 (0.92-1.32) | 0.276 |

### Adequate logistic regression model

Using Minitab software, we compute the binary logistic regression after the regression, confusion, and interaction assumptions. The probabilities are greater than 5%: thus, this model is adequate.

### Linear regression model: highlighted how was the exclusive effect of severe malaria

According to our statistical results, we have retained only 2 variables: response variable (Y) = Initial weight-Weight at 12 months; Predictive variable 1 (X_1_) = diagnosis of severe malaria. Finally, the equation retained was: Y = a + b_1_X_1_

### The Retained Predictor variable

Table 5 shows the retained predictive variable. We made our decision at an α = 0.05 level of significance i.e. if the p-value is <0.05, we reject the null hypothesis and otherwise we keep it. Since p is still very weak (p = 0.00005), the conclusion of a positive linear relation was even be declared very strongly.

### Retained predictor variable: severe malaria

The regression equation was: weight loss at 12 months = 15.3 + 0.697 diagnosis of severe malaria. We can use it for estimation purposes: we found y = 15.3 + 0.697X i.e. through linear regression the weight at the admission and 12 months was correlated with the effect of severe malaria.

### Interpreting the Linear Regression Model after consulting health data

The interpretation of the model is as follows: for each episode of severe malaria, the weekly averages of weight loss are in the order of 0.697 kg. Based on these forecasts of a weekly weight loss of 0.697 kg with a severe malaria episode, clinicians need to think what to suggest compensating for this, for us we suggest the maize, sorghum and soy.

There are correlations between the weight loss at 12 months under ART and the diagnosis of severe malaria on admission. R2 = 61.7% i.e.> 50%: this means that the variables explain the model at 61.7%. So, the model is good. The equation is a good predictor.

### Correlation between weight at 12 months on ART and severe malaria at admission

Table 6 showed the correlation between weight at 12 months on ART and severe malaria on admission. We see how the constant is 15,288 and the slope b is = 0.6973, this is the coefficient of severe malaria. In this case, because our p-value is <0.05 (Table 4), we confirm that there is a correlation between weight at 12 months on ART and severe malaria on admission. And how to say that we have a good model for the prediction? It is by the coefficient of determination R^2^. The coefficient of determination R^2^ = 0.617. Thus, only about 62% of the total variability in weight loss at 12 months under ART in the sample is explained by the linear regression relationship following energy expenditure because of severe malaria on the weight.

**Table 4.** Adequate logistic regression model.

| Method | Chi-square | DF | P |
| --- | --- | --- | --- |
| Pearson | 28.9468 | 26 | 0.314 |
| Deviance | 37.9923 | 26 | 0.061 |
| Hosmer-Lemeshow | 7.8017 | 8 | 0.453 |

**Table 5.** Retained predictor variable.

| Predictor | P | Null Hyp | Decision | Conclusion |
| --- | --- | --- | --- | --- |
| Constant | 0.011 | $\beta_0=0$ | Retain $H_0$ | The constant appears to be zero. Even so, we leave it in the model |
| Severe malaria | 0.00005 | $\beta_1=0$ | Reject $H_0$ | Severe malaria infection at admission can significantly contribute to explain the weight lost at 12 months |

**Table 6.** Analysis of table of coefficients of linear regression.

| Predictor | Coef | SE Coef | T | P |
| --- | --- | --- | --- | --- |
| Constant | 15.288 | 5.624 | 2.72 | 0.011 |
| Severe malaria infection | 0.6973 | 0.1003 | 6.95 | 0.00005 |

This example illustrates that there is no contradiction in finding other variables that contribute to weight loss called predictive variables Xi, i.e.: X_1_ = severe malaria; X_2_ = Weight on admission; X_3_ = CD_4_ / μl; X_4_ = co-infection HIV / severe malaria, X i + j = (5) diabetes, (6) cirrhosis, (7) tuberculosis, (8) cancer, (9) age, (10) poverty, (11) poor nutritional status, (12) helminths, etc.

From 9, 10, 11 predictive variables, there is really a conceptual problem to include all of them in the model.

### Consequence of such situation

One consequence of such a situation may be that clinicians should prevent episodes of malaria in HIV + patients living in malaria endemic areas.

### Analysis of variance for the four moments of weight measurement: 0, 3, 6 and 12 months

ANOVA was used to check for significant differences in variables and in different time periods. The significant level was set at p <0.05. Table 7 shows that the probability (0.591) is greater than 0.05; there is therefore no significant difference between the average of four weights. Using ANOVA, we find that the four means of weight were not significantly different.

**Table 7.** Analysis of variance for the four moments of weight measurement: 0, 3, 6 and 12 months.

| Source | DF | SS | MS | F | P |
| --- | --- | --- | --- | --- | --- |
| Regression | 3 | 129.15 | 43.05 | 0.64 | 0.591 |
| Residual Error | 124 | 8350.46 | 67.34 |  |  |
| Total | 127 | 8479.61 |  |  |  |

### Confounding factors

Not evaluated: The role of poverty and bad nutritional status as confounding factors, the role of other dysimmunities comorbidities (diabetic, cirrhose…).

### 83% of patients have not seen their weight increase

This is what was observed in the health data consulted, which underlines the need for a supplement and other antimalarial measures, lifestyle, diet following the second specific objective.

### Repetition of malaria episodes based on the number of CD4 /μl at admission

In sum, there is no statistically significant interaction of the diagnosis of severe malaria on the number of CD_4_ <50 cells / μl on admission, although it has been observed in some individual cases.

### The effect of clinical malaria infection on weight

Based on our example of 72 observations, we make predictions of the effect of clinical malaria infection on weight.

We identified more than 7 episodes of severe malaria (30%) and a CD_4_ count <50 cells / μl in the subgroup whose weight did not increase in the first year under ART.

The other subgroup whose subjects had gained weight in 12 months under ART had 5% of severe malaria attacks and a CD_4_ count> 50 cells /μl. CD_4_ with less than 7 malaria episodes per year.

HIV infection increases the repetition of clinical malaria episodes, which could be associated with weight loss.

Our results emphasized that the accurate assessment of the effect of clinical malaria on HIV-infected people is limited by the lack of rigorous diagnostic criteria and the definition of what may be considered malaria.

If the thick-film had been made and was a test with the highest sensitivity-specificity and number of parasites / μl of blood, patients could have a parasitaemia that coincides with the fever of another origin (such as opportunistic infections, bacterial diseases such as *Streptococcus pneumoniae*, *Salmonella typhi* species or not).

## Discussion

### Key results with reference to study objectives

The health data from a sample of 72 participants of AMOCONGO Kinshasa Kasavubu HIV Register show overall, in the whole group (including the HIV + subgroup whose weight did not increase in the first year on ART and the subgroup who gained weight), the ANOVA admission, at 3 months, at 6 months and at 12 months later the four weight averages are not significantly different. So, there is on average no significant weight change in the first year under ART. We conclude that severe malaria is the cause of weight loss and should be controlled by the preventive treatment of malaria. Whence continuous treatment of malaria in a HIV positive subject (with therapeutic antimalarials intermittent treatment) will help prevent weight loss by decreasing the parasite biomass buried in the deep organs (liver, spleen, brain, kidney) therefore not detectable at the peripheral level with the examination of the thick drop.

### Outcomes of the study

The present study aimed to determine the relationship between severe malaria and HIV among HIV+ adults living in malaria endemic area as Kinshasa in the Democratic Republic of the Congo and their clinic expression in the weight loss. About malaria definition based on clinical signs, this study is in conformity with Flateau [3] and Rogier [32]: Flateau affirms that because of the absence of malaria rigorous diagnostic criteria, the precise evaluation of the effect of malaria in HIV-infected patients is limited [3]. Rogier said that it is difficult to define malaria although its epidemiological data are known [32]. For us the rigorous diagnostic criteria are a positive microscopic test before treatment and the disappearance of admissions symptoms after the AMT as we stated that malaria is a retrospective diagnosis using health data. It means that when a person presents malaria symptoms you never know if it’s malaria or not. It is only after the treatment that you’ll get the answer because of co-infections.

According to the malaria management our study agrees to wait for microscopy results before starting the treatment. We found for 10 years (2000-2009) only 32% of malaria positives samples in a study that we evaluated in Kinshasa University Hospital about malaria microscopic diagnosis [35]. For us in area of high endemicity of malaria, a treatment should begin with a serious sign as fever (39-40°C), positive microscopic test and anemia that may be malaria for an adult with HIV living in high malaria endemicity area. Although if the microscopic test is negative, the fever can have another origin such as bacterial diseases particularly tuberculosis, *Streptococcus pneumonia*, non-typhi Salmonella species and other opportunistic diseases.

As the malaria mortality is increasing with the severity of immunosuppression (low CD4 cells / μl) [3], this study suggests treating malaria if CD_4_4<50 cells / μl in malaria area with clinical signs and malaria microscopic positive test according to our observation of the health data. We chose our sample based on physician’s diagnosis made with severe malaria symptoms as fever and anemia.

### Malaria episodes and HIV infection

The present study identified more than 7 episodes (30%) of severe malaria with a number of CD_4_4<50 cells/μl in the sub-group who lost weight in the first year of ART. The other subgroup who gains weight had 5% of 1 to 7 malaria episodes. In Nigeria Amuta observed that in people living with HIV and AIDS, the prevalence of malaria was low when CD_4_ counts increased and it was increased when CD_4_ was low [31]. In a prospective study in South Africa, HIV-infected people had an increased risk of severe malaria by 4 times, and the prevalence of severe malaria was the highest when the CD_4_ cell count was less than 200 cells per μl [3]. In subgroup analyses, HIV infection was associated with an increased risk of severe malaria in non-immune, but not in semi-immune people [3].

### Conceptual Model ◻ on the associations between malaria, HIV and weight changes

The conceptual model explains the physiopathology of the co-infection HIV-MALARIA. In one hand malaria stimulate the NADPHase blocked by the ART making that a HIV+ people under ART can lose weight during malaria crisis. In the other hand the key of that physiopathology is relative to the endothelium equilibrium to establish between oxidants and anti-oxidants factors as selenium in the cellular level.

### Therapeutic nutrition

In view of the above, we recommend that therapeutic nutrition be included in the overall strategy to combat HIV-*Plasmodium* co-infection.

## Conclusions

Based on these forecasts of a weekly weight loss of 0.697 kg with a severe malaria episode, we suggest the consideration of the NADPH oxidase in the physiopathology of the co-infection HIV-malaria for therapeutic relevance using local foods rich in Selenium. Excluding other variables, malaria is the main cause of weight loss under ART in Kinshasa. We found that health data that reports longitudinal data adhere to generally accepted Prospective Study definition.

## Acknowledgments

None.

